# A Molecular Image-based Explainable Artificial intelligence Framework Identifies Novel Candidate Antibiotics

**DOI:** 10.1101/2024.07.21.604494

**Authors:** Kingsten Lin, Yuxin Yang, Feixiong Cheng

## Abstract

The continued growth of antibiotic resistance and the slowing of antibiotic discovery poses a large challenge in fighting infectious diseases. Recent advances in Artificial intelligence (AI) technologies offer a time- and cost-effective solution for the rapid development of effective antibiotics. In this study, we presented an explainable AI framework from a pre-trained model using 10 million drug-like molecular images. Specifically, we created a fine-tuned ImageMol from experimental *Staphylococcus aureus* inhibition assays which contained 24,521 molecules consisting of 516 active compounds and 24,005 non-active compounds. Our optimized AI model achieved a strong AUROC of 0.926. The model was then used to predict the antibiotic activities from 10,247 FDA-approved, clinically investigational, or experimental molecules from the DrugBank database. After further filtering, 340 molecules were identified to have antibacterial behavior while simultaneously being dissimilar to known antibiotics. Finally, 76 candidates were identified as FDA-approved drugs for other applications. Thus, those candidates can be repurposed into needed novel antibiotics. We further illustrated explainable molecular images for top predicted candidate drugs via Gradient-weighted Class Activation Mapping (Grad-CAM) heatmap analysis. In summary, the presented molecular image-based AI model in drug discovery could be highly favorable due to its high performance, speed, and biological interpretation.

## Introduction

The antibiotic resistance crisis has been at the forefront of medicine with resistance rapidly evolving (Alós, 2015) and the discovery of new antibiotics slowing (Wohlleban, 2016). One such example of a problematic bacteria is *Staphylococcus aureus*, a gram-positive bacteria that causes many severe diseases including but not limited to sepsis, dermatitis, osteomyelitis, and endocarditis. Methicillin-resistant *Staphylococcus aureus* (MRSA) has become widespread, especially in medical settings, leading to many hospital outbreaks (Lee et al., 2018). The β-lactam resistance of MRSA makes it especially difficult to treat with common antibiotics such as penicillins. It is urgently needed to develop novel and effective antibiotics.

With the recent advancements in Artificial intelligence (AI) technologies, though, antibiotic discovery can be aided through *in silico* predictions that can quickly work through large chemical databases that otherwise would have been very difficult to screen experimentally (Zeng et al., 2022a; Zhou et al., 2020b; Cheng et al., 2024). For example, the novel, broad-spectrum antibiotic halicin was discovered from a database of more than 6,000 compounds in Stokes et al. (2020). Indeed, AI models offer effective tools for drug development (Zeng et al., 2022a; Fang et al., 2020; Cheng et al. 2024; Qiu & Cheng, 2024), including antibiotics discovery (Stokes et al., 2020; Wong et al., 2024).

Traditional AI models, however, must address many challenges. One of those is their black-box nature. The complexity of AI models means their predictions often seem to follow no explainable rationale, which poses a problem in biologically interpreting AI predictions (London, 2019). This is especially crucial in antibiotic prediction, where antibiotics often come in classes of similar structures and mechanisms. If an AI model for predicting antibiotics were interpretable, entire classes of antibiotics could be discovered instead of lone compounds, thus greatly increasing the efficiency of drug exploration (Wong et al., 2023).

Additionally, machine learning for drug design must always consider how to best represent chemicals (Li et al, 2021). Many approaches have been used, including various sorts of molecular fingerprints (Muegge, 2015), as well as automatic representations such as graphs (Stokes et al., 2020) or simplified molecular-input line-entry system (SMILES) strings (Chithranada, 2020). Recently, improvements in computer vision have motivated the usage of image-based molecular representations. Images do not require the feature extraction process that fingerprints do and can be more accurate than string representations. Additionally, images are more interpretable than strings or fingerprints. ImageMol, a molecular image-based AI framework, was previously used to predict anti-SARS-CoV-2 inhibitors during the COVID-19 pandemic and successfully identified anti-SARS-CoV-2 drugs (Zeng et al, 2022b). Applying the framework to other problems such as antibiotic discovery naturally follows.

In this study, we further utilized ImageMol to predict the *S. aureus* inhibition ability of 10,237 molecules from the DrugBank database (Knox, 2024). After, the resulting 3,650 predicted hits were further filtered by a high confidence threshold of 0.95. The result was 794 strong candidates, 340 of which were structurally dissimilar to existing antibiotics, and 76 of which were additionally FDA-approved (Figure 1). Thus, those 76 molecules can now be investigated further, potentially leading to the discovery of a novel antibiotic. Indeed, example molecules from the final dataset were demonstrated to hold promise through analysis of the literature. Additionally, latent space representation of the DrugBank dataset suggested that the model understood antibiotic classes. This idea was further supported by Gradient-weighted Class Activation Mapping (Grad-CAM) analysis (Selvaraju et al., 2019) that showed how ImageMol focused on specific substructures. Therefore, image-based frameworks such as ImageMol can not only efficiently make predictions but also be used in an explainable way. By understanding the model, useful patterns and information can be extracted. For example, important substructures can be identified and leveraged to make *de novo* antibiotics.

**Figure 1:**
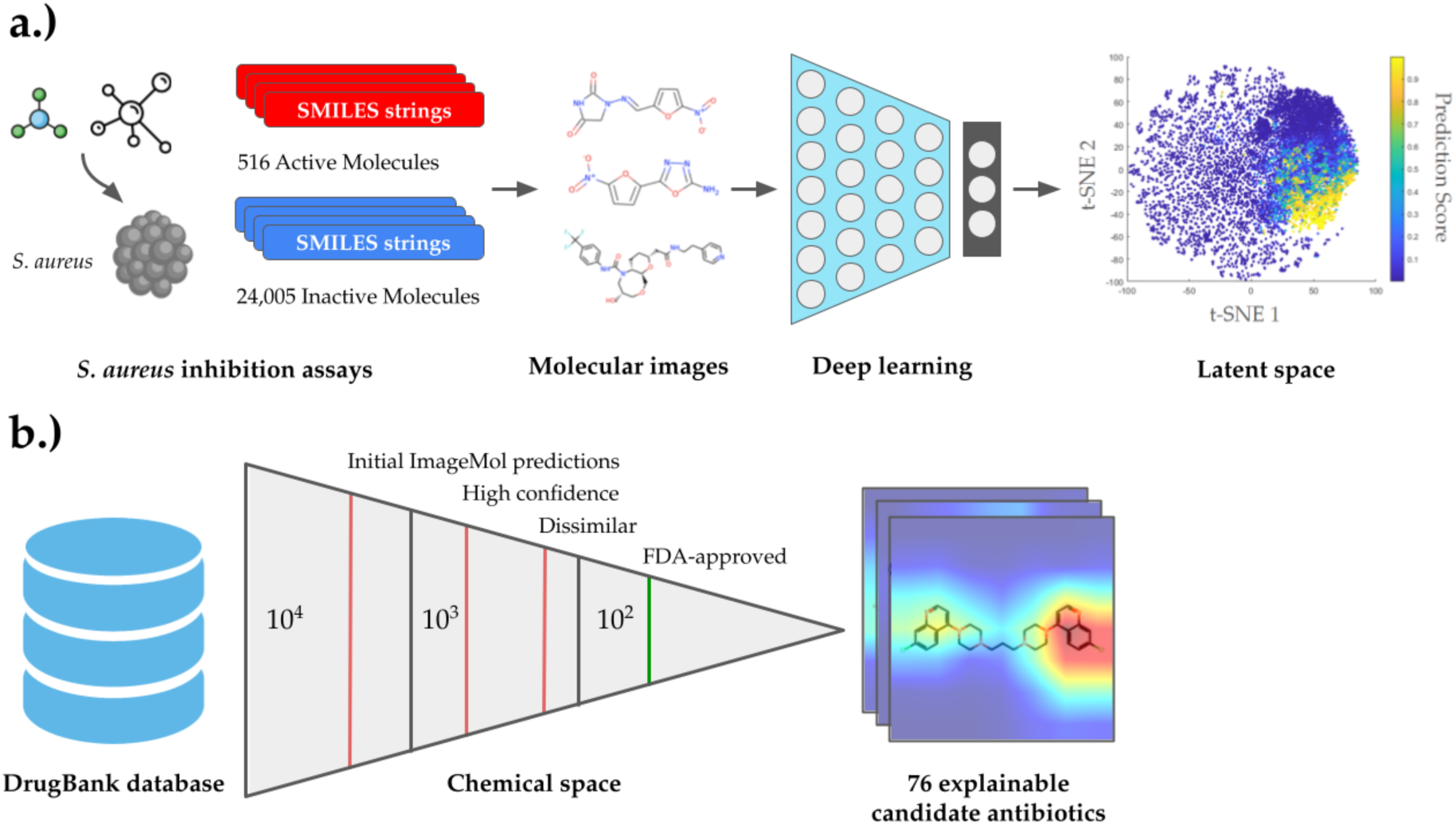
Graphical paper description. **a.)** Description of the fine-tuning process. SMILES strings were converted into molecular images and fed into the ImageMol framework. A final layer (black) was then appended to the pretrained molecular encoder (blue). The latent feature space was then visualized using sklearn’s t-SNE with default parameters. **b.)** Description of the molecular workflow. Over 10,000 molecules from the DrugBank are screened by the ImageMol model and filtered for low Tanimoto similarity, high confidence, and FDA approval. The results include 76 candidate antibiotics. The rationale for antibiotic prediction can be visualized using Gradient-weighted Class Activation Mapping (Grad-CAM) heatmap analysis.

## Results

### Fine-tuning to predict antibiotic profiles

To make predictions on antibacterial profiles, the pretrained ImageMol model (*cf.* Methods) was fine-tuned (**Figure 1**) on a binarized assay measuring *S. aureus* inhibition from Wong et al. (2024). The assay contained 24,521 molecules, 516 of which were considered active by a 20% relative to mean growth cutoff (**Supplementary Table 1**). Thus, 24,005 molecules were inactive. To fine-tune, the original SMILES strings from the assay were canonized and converted into two-dimensional image representations. Then, ImageMol was trained by appending a fully connected layer to the pretrained model and minimizing cross-entropy loss between class probability and true labels. This training was completed using the fine-tuning script included with the ImageMol framework. To increase performance, hyperparameter tuning methods such as Bayesian optimization were explored. Ultimately, though, ImageMol default hyperparameters resulted in the best performance and thus were used (*cf.* Methods). The resulting fine-tuned model was then evaluated using an external test set, where it achieved an impressive AUROC score of 0.926 (**Figure 2**). ImageMol’s strong performance is further shown by a high recall value of 0.92 (**Table 1**).

**Figure 2:**
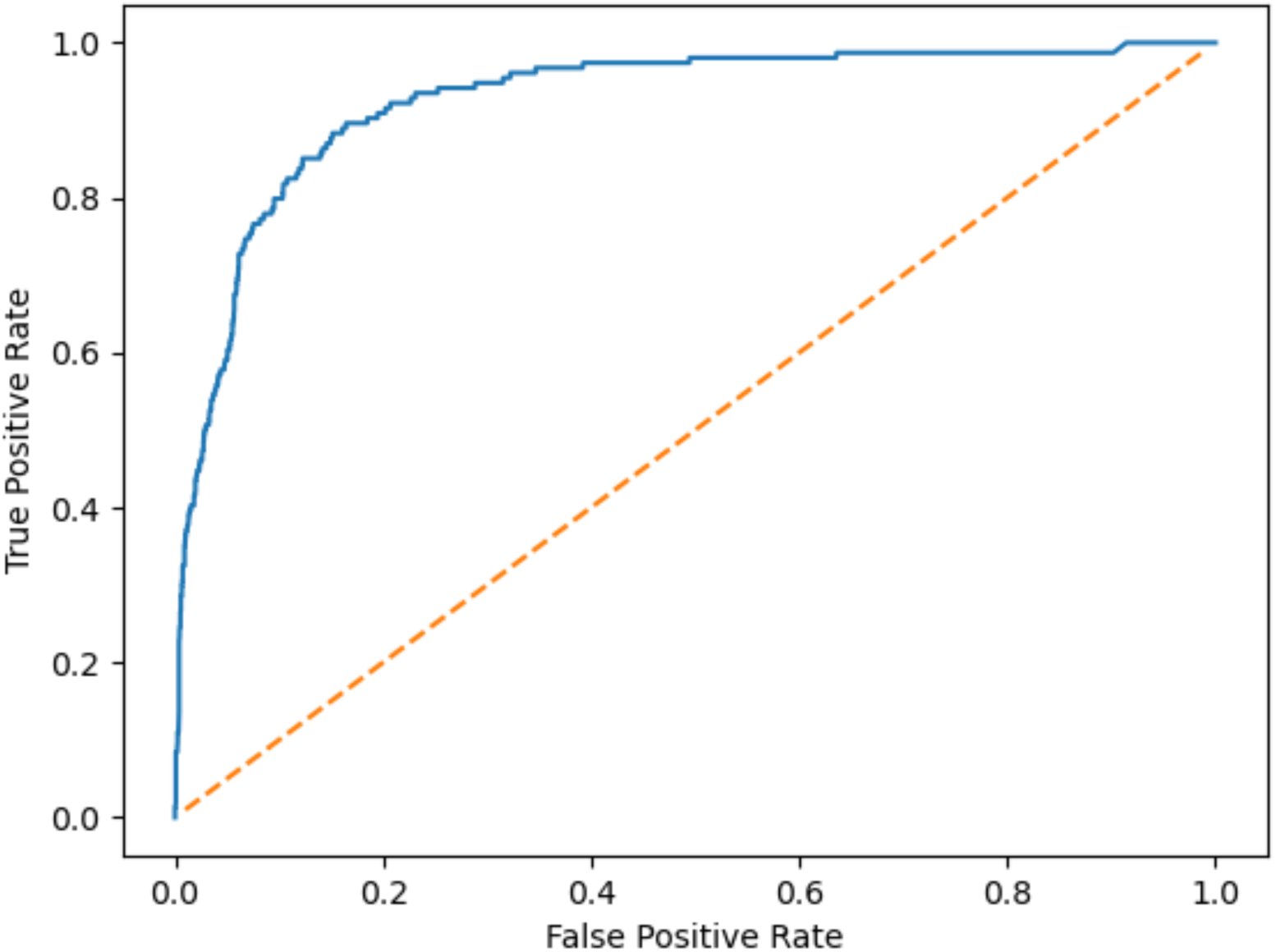
Receiver Operator Characteristic (ROC) curve for *S. aureus-*fine-tuned ImageMol. AUC = 0.926, calculated using a trapezoidal Riemann sum. The curve was made using an unseen test dataset.

**Table 1:** Confusion matrix for *S. aureus-*fine-tuned ImageMol.

|  |  |  |  |
| --- | --- | --- | --- |
| Recall = 0.92 | Predicted Positive | Predicted Negative | Total |
| Actual Positive | 142 | 12 | 154 |
| Actual Negative | 514 | 1,785 | 2,299 |
| Total | 656 | 1,797 | 2,453 |

### Virtual screening of antibacterial activities from DrugBank

The fine-tuned model was then used to predict the *S. aureus* inhibitive ability of 10,247 molecules from the DrugBank database, of which included 1,974 FDA-approved, 2,649 clinically investigational, and 5,624 experimental or withdrawn compounds. The DrugBank database was chosen for its utility in drug repurposing, as drugs which are already approved for other purposes would have an easier and faster approach to be approved as much-needed novel indications (Zhou et al., 2020a; Fang et al., 2021; Zhou et al., 2020b), such as antibiotics.

The initial predictions resulted in 3,650 hits, which were defined as a prediction score of over 0.5 (**Supplementary Table 2**). Because the DrugBank dataset includes known antibiotics, those antibiotics can be used as an internal control to validate the model. A simple name-based screening method was used to identify antibiotic compounds throughout the entire dataset. The DrugBank database included the names of each compound, and so those names were checked for strings associated with antibiotics such as “cef” from cephalosporins, “cillin” from the penicillin class, and “cycline” from tetracyclines (**Supplementary Table 3**), resulting in a list of 153 molecules. The model labeled 97% of those known antibiotics as being active against *S. Aureus*. Upon manual inspection of the five mislabeled molecules, it was found that they all were not antibiotics. Thus, all of the known antibiotics identified by the name screening were correctly predicted to show S. aureus inhibition, suggesting the validity of the predictions. Additionally, more than 80% of the 148 antibiotics were predicted to be active with a probability of over 0.95, suggesting that during further filtering, only keeping molecules with a score greater than 0.95 will lose few antibiotics while greatly reducing the search space. Of note, 86 of the 148 antibiotics (58%) had not been seen by the model during fine-tuning, suggesting that this strong performance is not the product of an overfit. Thus, this metric demonstrates the validity of the model, as it does not miss known antibiotics.

Using t-SNE to represent the latent space of the model, specific classes of antibiotics can be seen clustered together (**Figure 3**). This suggests that the fine-tuned ImageMol can identify classes, an important feature for interpretability. Grad-CAM heatmap analysis can additionally be used to visualize the rationale behind the model. Once more, the model’s ability to identify class-wide features can be seen. For example, in **Figure 4**, the heaviest attention is put on the signature β-lactam-thiazolidine ring in each penicillin. ImageMol’s ability to identify classes and substructures is especially exciting due to the usual black-box nature of ML models. By identifying substructures associated with antibiotic behavior, *de novo* antibiotics can be explored by building off of those substructures, potentially leading to an entire novel class of antibiotics.

**Figure 3:**
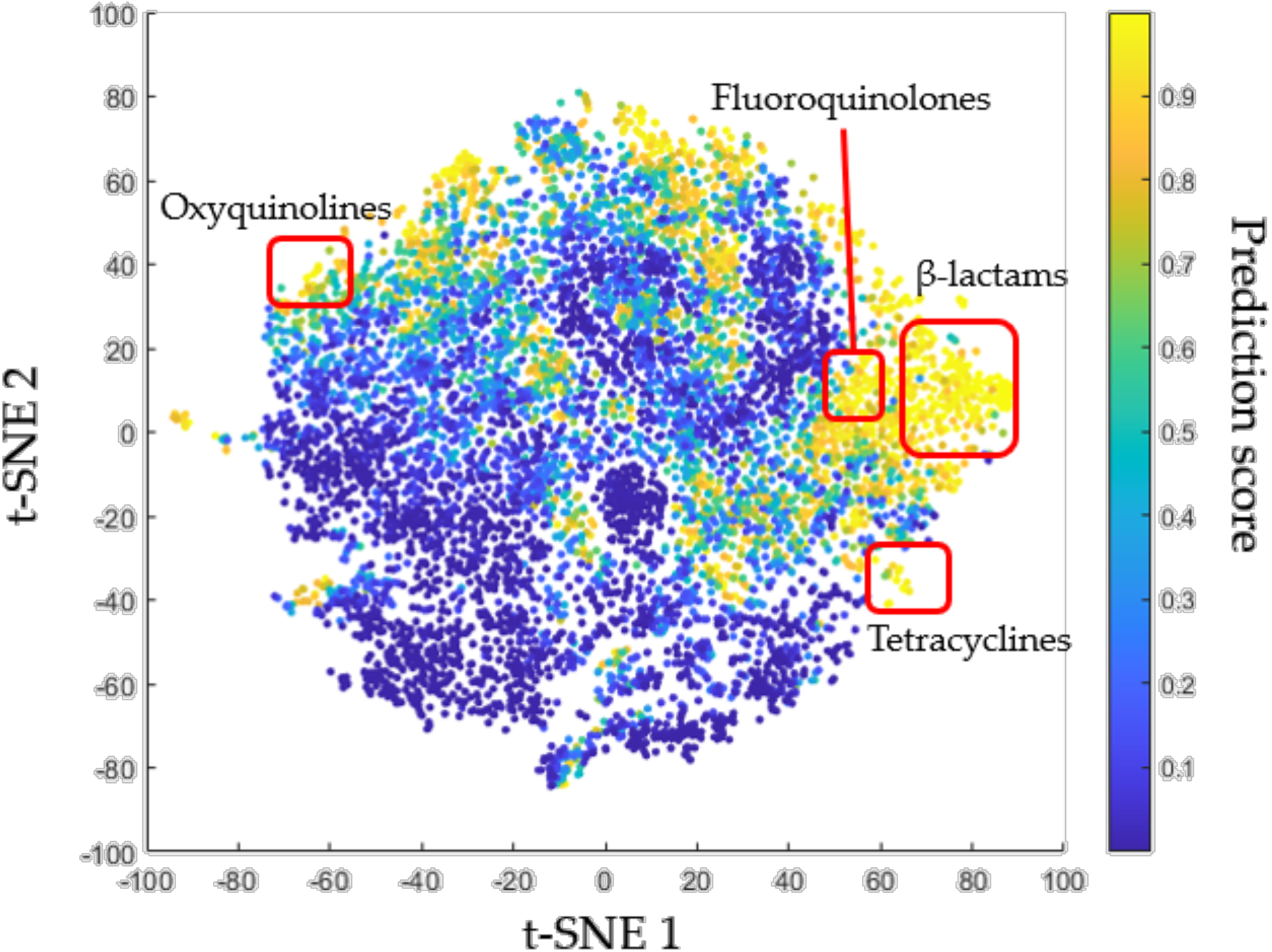
t-SNE visualization of 10,237 molecules from DrugBank dataset, colored by antibacterial behavior prediction score. Stronger yellow means the AI identified the molecule as more likely to be an antibiotic. Selected antibiotic classes are labeled. t-SNE analysis was completed using sklearn’s TSNE (Pedregosa et al., 2011) with default parameters.

**Figure 4:**
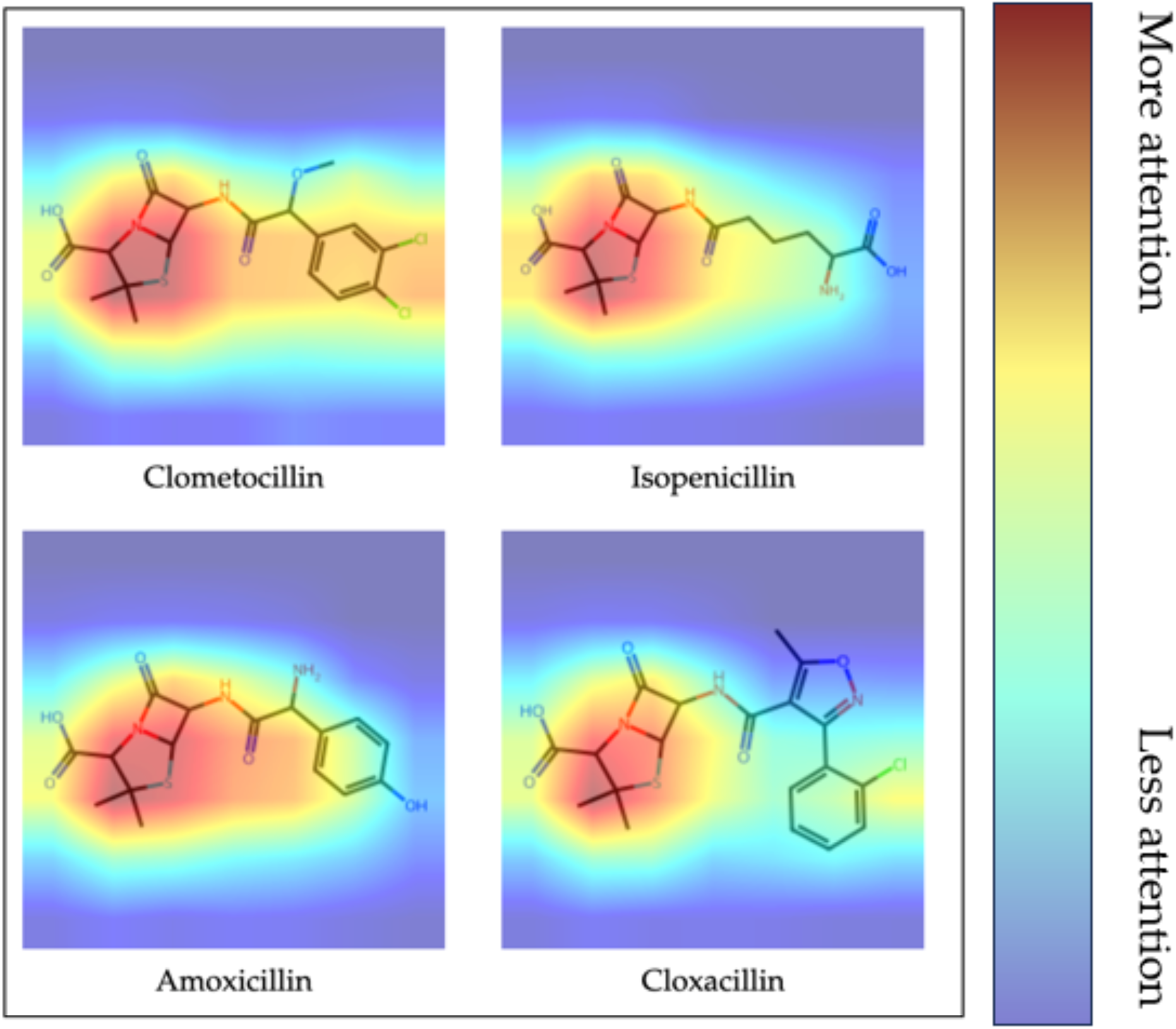
Grad-CAM analysis of four penicillin-class molecules, demonstrating substructure awareness. Warmer colors indicate higher ImageMol attention. In all four of these molecules, the model appropriately places the most weight on the β-lactam-thiazolidine ring, the defining feature of penicillins.

### Identifying potential antibiotics from FDA-approved drugs

After the initial fine-tuned model predictions, the molecules were further filtered. First, only molecules with a score of greater than 0.95 were kept, resulting in 794 molecules. To ensure that only novel molecules were kept, a max Tanimoto similarity metric was used. Using RDKit, each molecule was compared to the 148 antibiotics identified in the earlier name screen. Then, the largest Tanimoto similarity value was recorded. Molecules with a max similarity of less than 0.4 were kept, resulting in a set of 340 molecules. Grad-CAM heatmap analysis was performed on the 340 high-confidence and dissimilar molecules, and notable substructures were recorded. Some of those notable substructures can be seen in **Figure 5**. The model’s ability to consistently identify and put a high weight on these substructures implies an understanding of those substructures and suggests their chemical importance. Additionally, experimental validation of predicted candidate compounds is highly warranted in the future.

**Figure 5:**
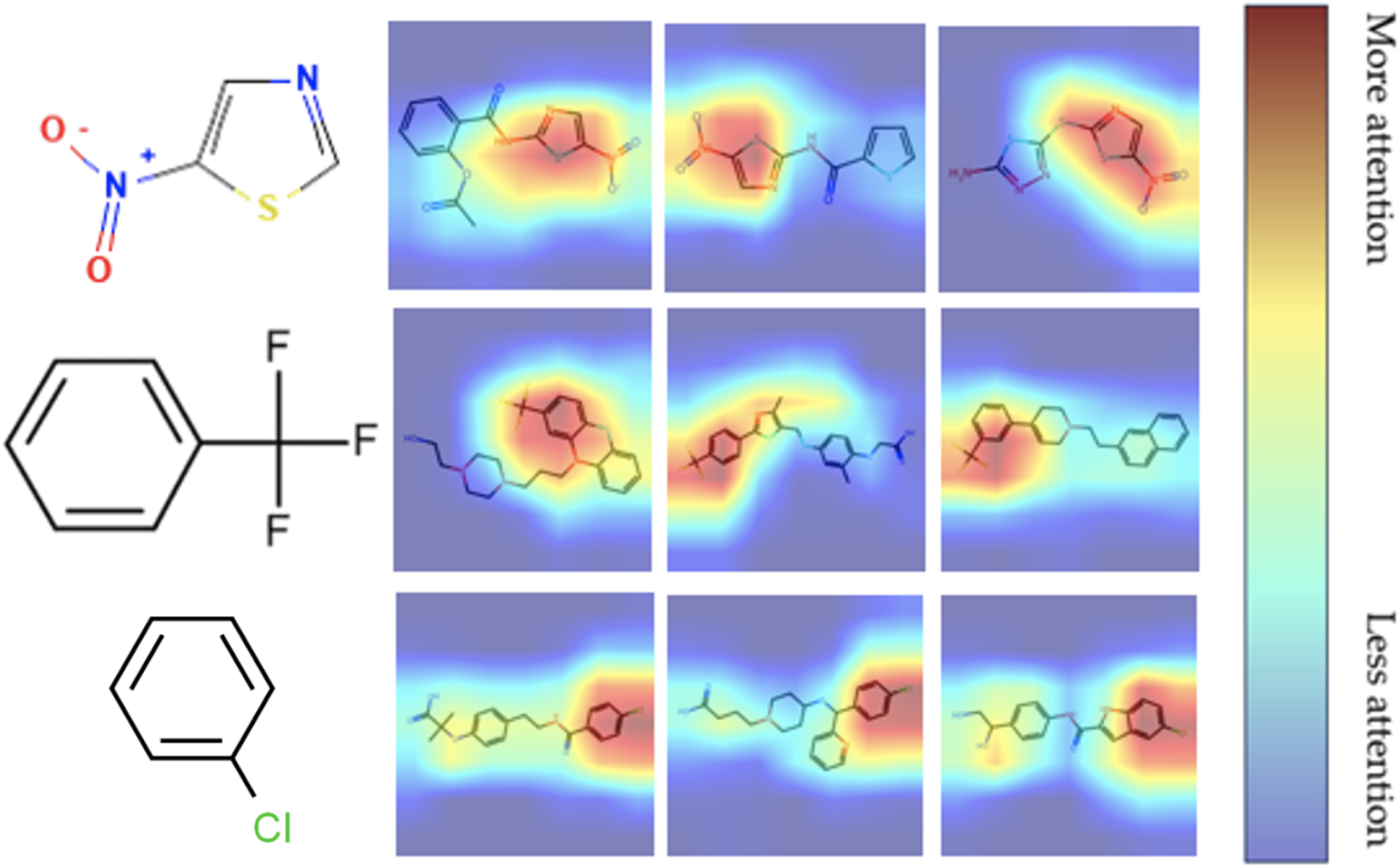
Selected molecules with highly weighted substructures from Grad-CAM analysis. Warmer colors indicate higher ImageMol attention. On the right are Grad-CAM heatmaps of various molecules from the set of 340 high-confidence and dissimilar molecules. On the left are three example substructures that can be seen in the example heatmaps.

To produce the final set of molecules, FDA-approved molecules were identified and kept. The final results included 76 molecules (**Table 2 and Supplementary Table 4**). Literature review of various molecules from that set suggests their promise as antibiotics. For example, the antifungal oxiconazole had strong activity against *S. aureus in vitro* in Kaul et al. (2023). Fluphenazine, an antipsychotic with a probability of 0.961, was recorded having broad-spectrum antibiotic behavior by Dastidar et. al (1995). Nitazoxanide, traditionally used for treating diarrhea-causing infections, was also identified by Kaul et al. (2022) as a potential answer to *S. aureus*. These three drugs represent molecules that are in the final prediction set and are directly supported by studies. Additionally, there are molecules in the final set that have not been directly investigated but still have potential. For example, Siponimod is a sphingosine-1-phosphate (S1P) receptor modulator that was developed for Multiple Sclerosis. Zore et al. (2022) found that a different S1P receptor modulator named Etrasimod had great potential as an antibiotic targeting gram-positive bacteria, potentially suggesting the viability of Siponimod as well. Another molecule is the antimalarial drug Piperaquine, which is suspected to inhibit glutathione S-transferase (GAT). Previously, Pugazhendhi et al. (2017) suggested that GST is important to antibacterial drug resistance, implying that GST inhibition could be vital to treating tough bacteria like MRSA. These two drugs are representative of molecules that have not been explored as antibiotics but are rational candidates. All five example drugs presented here, as well as the drugs in the final dataset, represent exciting candidates for further study and have the potential to become novel antibiotics. Grad-CAM analysis of those five molecules again identifies specific substructures (**Figure 6**). Further study of these substructures can further inform the drug discovery process, including the production of *de novo* candidates.

**Figure 6:**
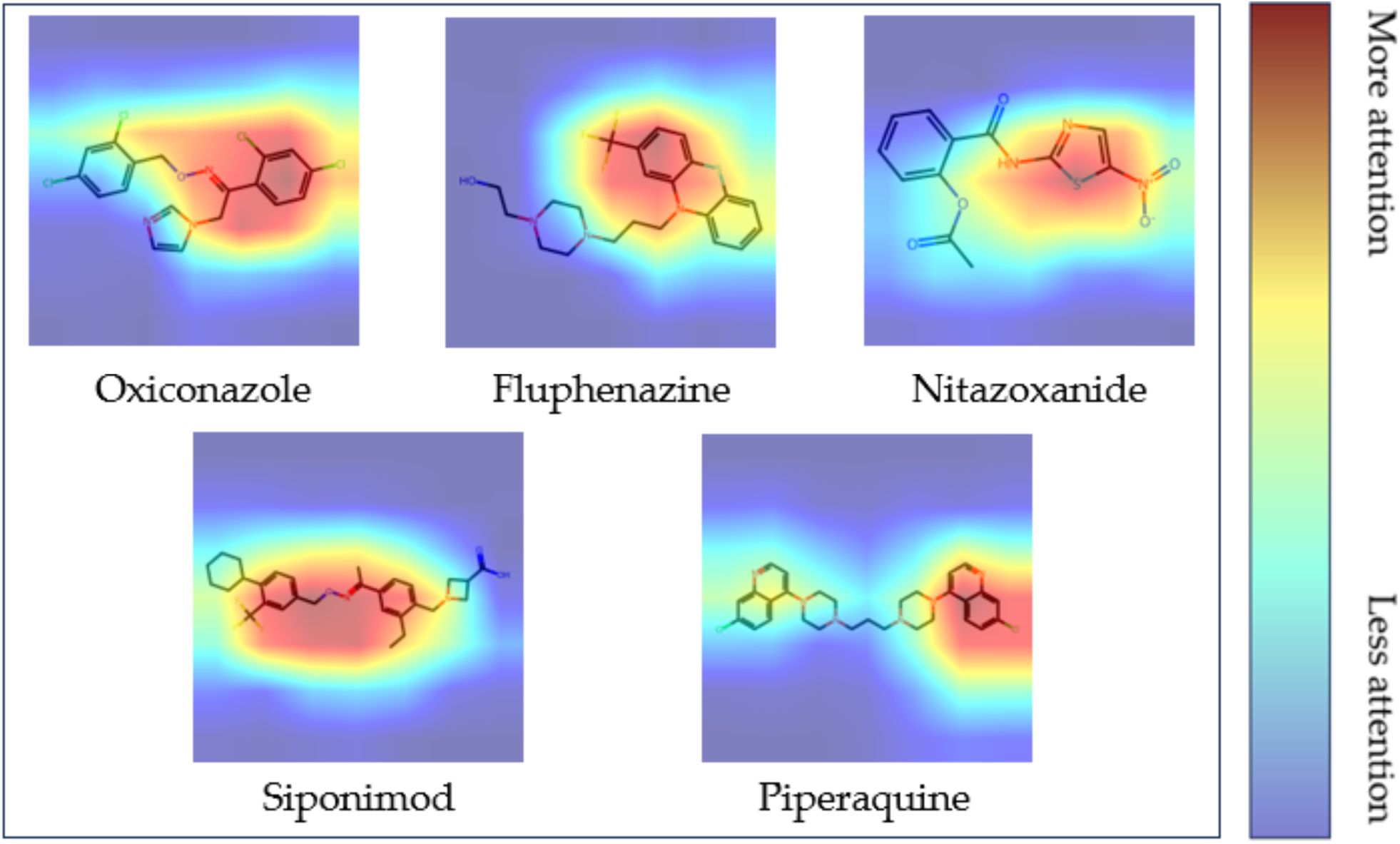
Grad-CAM analysis of five example molecules from the final prediction set. Warmer colors indicate higher ImageMol attention. Of note, distinct substructures can be seen and interpreted. For example, there is particularly high weight placed on the trifluorobenzene in Fluphenazine, or on the chlorobenzene and pyridine in Piperaquine.

**Table 2:** Ten top performing molecules from the final filtered set.

| Name | MaxSimilarity | Probability |
| --- | --- | --- |
| Oxiconazole | 0.32777778 | 0.9996221 |
| Acitretin | 0.2551382 | 0.9986223 |
| Niclosamide | 0.3175583 | 0.998302 |
| Dequalinium | 0.39236393 | 0.9982206 |
| Fusidic acid | 0.36321304 | 0.9976229 |
| Isoconazole | 0.30769231 | 0.9968772 |
| Bronopol | 0.0816092 | 0.9962607 |
| Piperaquine | 0.38931298 | 0.99606365 |
| Tafamidis | 0.37323511 | 0.99583036 |
| Nitisinone | 0.39643268 | 0.9956558 |

## Discussion

ImageMol is an image-based ML framework that benefits from high performance (Figure 2) and low resource cost. In this study, ImageMol was leveraged to predict 3,650 potential antibiotics, out of which 148 are established antibiotics and 76 are FDA-approved drugs that have promise in being repurposed into novel antibiotics. Further inspection of literature additionally revealed the viability of various molecules’ mechanisms of action. These findings are important to the search for novel antibiotics and represent excellent candidates for screening.

Additionally, the interpretability of ImageMol was demonstrated through Grad-CAM heatmap analysis, where important substructures can be easily identified (**Figures 4-6**). This paper shows the promise of a specific strategy for drug discovery using ML. With image-based frameworks such as ImageMol, predictions can be made quickly and analyzed intuitively through substructures. It is important to note that this explainability is made especially simple through the model’s image-based identity. The interpretability then lends itself to better human understanding of antibiotic mechanisms. An interesting possibility with this development is using ImageMol and Grad-CAM analysis on existing antibiotic datasets to potentially extract further information regarding known antibiotics. A better understanding of known antibiotics can then lead to the more informed discovery of novel antibiotics.

Some limitations must be acknowledged. First, the Grad-CAM analysis could benefit from a numerical metric. The methods in this study used human inspection to identify strongly weighted substructures, but an automated method would be faster and more objective. Such a method could consider the area of attention on non-white space in pixels, or perhaps even use a separate computer vision model to identify substructures. Secondly, it must be noted that this entire study is contained *in silico*. Thus, *in vitro* screening is still needed to confirm the predictions. This is one of the natural next steps of this paper. The other next step is to continue to expand the substructure-based process outlined in this study. A basic pipeline has been demonstrated: fine-tune the model, analyze the results with Grad-CAM, and identify substructures. The process can be continued by using those substructures to create *de novo* compounds, and then testing those new molecules for antibiotic behavior in hopes of identifying many novel antibiotics off of just one hit. Explainable antibiotic discovery facilitated through image-based machine learning offers an exciting way to both better understand and better predict antibiotics.

## Materials and Methods

### ImageMol framework

ImageMol, by default, runs on the ResNet-18 architecture and uses 224 x 224 pixel images as inputs. The model pretraining includes five tasks: multigranularity chemical cluster classification, molecular rationality discrimination, jigsaw puzzle prediction, mask-based contrastive learning, and molecular reconstruction. These tasks aim to ensure that the model can, respectfully, recognize chemical substructures, identify if a molecule was chemically possible, recognize shuffled images, identify images with small sections that are masked out, and remake molecular images from latent features. The loss associated with each task is summed to a total loss, which is then optimized with mini-batch stochastic gradient descent in the pretraining process. In total, these tasks give ImageMol a better general understanding of chemicals. Additionally, the pretraining dataset is augmented with grayscale, rotation, and reflection to further induce generalization. Fine-tuning is completed by appending another full connection layer to the model and optimizing cross-entropy loss between the class probability and the true label. Zeng et al. (2022b) contains the full explanation of the ImageMol framework.

### Fine-tuning dataset

The fine-tuning dataset was obtained from Wong et al. (2023). Their method first dissolved compounds in dimethyl sulfoxide at 5mM concentration. Then, *S. aureus* was grown in Luria broth (LB) medium and diluted 1:10,000 to be put in multi-well plates. The dissolved compounds were finally added to the plates containing *S. aureus* to achieve a 50 µM concentration. After incubation at 37°C, growth was quantified using a SpectraMax M3 plate reader and normalized to the interquartile mean of each plate. The method is described in full in Wong et al. (2023).

### Fine-tuning and running ImageMol

The *S. aureus* inhibition assay was binarized using a 20% relative to the mean cutoff, resulting in 516 active molecules out of 24,521 total. Then, the SMILES strings from the original assay were canonized and converted to images using the utility scripts included with ImageMol. An Optuna (Akiba et al., 2019) study to optimize hyperparameters was attempted. However, time constraints required that the study only operate in a limited capacity, with a smaller number of epochs. As a result, the default parameters achieved better performance compared to the Optuna-defined parameters. Thus the final fine-tuned model used a learning rate of 0.08, a weight_decay value of −5, and a momentum value of 0.9 as is standard in the ImageMol code. However, the model was configured to use stratified split and global imbalanced weight, due to the heavily imbalanced dataset.

To use ImageMol to make predictions, the DrugBank dataset had to be processed in a similar way to the *S. aureus* assay, with SMILES canonization and image creation. Then, the predictions were obtained by applying the fine-tuned model to the DrugBank dataset. Grad-CAM analysis was performed using the script included in the ImageMol package.

### Data analysis

In Figure 2, the AUC was calculated using a trapezoidal Riemann sum. The ROC itself utilized 10,000 thresholds. In Figure 3, the latent space was recorded by replacing the final layer of the fine-tuned ImageMol model with an identity later and then running the *S. aureus* assay through the model. Then, t-SNE analysis was performed with sklearn’s TSNE using default parameters.

### DrugBank database

The DrugBank database used for predictions was accessed through an academic license. Any researcher who wishes to replicate the results of this paper should also obtain the free academic license to gain access to DrugBank. The database is a curated library that stores the pharmaceutical data of thousands of molecules. The .csv file of DrugBank database structures was downloaded and processed using the SMILES canonization and image conversion utility scripts included with ImageMol. After canonization, the total dataset size was 10,247 molecules. That set was then used for predictions.

Using the “Drug Groups” column of the DrugBank database, a molecule’s FDA approval or investigational status could be checked. For this study, molecules that were approved and not withdrawn were considered as “approved” and molecules that were investigational and not approved were considered as “clinically investigational.”

## Author Contributions

F.C. conceived the study. K.L. performed all experiments and data analysis. Y.Y. interpreted the data analysis. K.L. draft the manuscript and both K.L. and F.C. critically revised the mansucript. All authors gave final approval of the manuscript.

## Availability of Code and Data

All code and pre-trained models are freely available at https://github.com/ChengF-Lab/imageMOL

## References

Akiba, T., Sano, S., Yanase, T., Ohta, T., & Koyama, M. (2019). Optuna: A Next-generation Hyperparameter Optimization Framework. arXiv. https://arxiv.org/abs/1907.10902

Alós J. I. (2015). Resistencia bacteriana a los antibióticos: una crisis global [Antibiotic resistance: A global crisis]. Enfermedades infecciosas y microbiologia clinica, 33(10), 692–699. 10.1016/j.eimc.2014.10.004

Boelrijk, J., Pirok, B., Ensing, B., & Forré, P. (2021). Bayesian optimization of comprehensive two-dimensional liquid chromatography separations. Journal of chromatography. A, 1659, 462628. 10.1016/j.chroma.2021.462628

Cheng, F., Wang, F., Tang, J., Zhou, Y., Fu, Z., Zhang, P., Haines, J. L., Leverenz, J. B., Gan, L., Hu, J., Rosen-Zvi, M., Pieper, A. A., & Cummings, J. (2024). Artificial intelligence and open science in discovery of disease-modifying medicines for Alzheimer’s disease. Cell Reports Medicine, 5(2), 101379. 10.1016/j.xcrm.2023.101379

Chithrananda, S., Grand, G., & Ramsundar, B. (2020, October 23). Chemberta: Large-scale self-supervised pretraining for molecular property prediction. arXiv.org. https://arxiv.org/abs/2010.09885

Dastidar, S. G., Chaudhury, A., Annadurai, S., Roy, S., Mookerjee, M., & Chakrabarty, A. N. (1995). In vitro and in vivo antimicrobial action of fluphenazine. *Journal of chemotherapy (Florence*, Italy), 7(3), 201–206. 10.1179/joc.1995.7.3.201

Fang, J., Zhang, P., Zhou, Y., Chiang, C. W., Tan, J., Hou, Y., Stauffer, S., Li, L., Pieper, A. A., Cummings, J., & Cheng, F. (2021). Endophenotype-based in silico network medicine discovery combined with insurance record data mining identifies sildenafil as a candidate drug for Alzheimer’s disease. Nature aging, 1(12), 1175–1188. 10.1038/s43587-021-00138-z

Fang, J., Zhang, P., Wang, Q., Chiang, C. W., Zhou, Y., Hou, Y., Xu, J., Chen, R., Zhang, B., Lewis, S. J., Leverenz, J. B., Pieper, A. A., Li, B., Li, L., Cummings, J., & Cheng, F. (2022). Artificial intelligence framework identifies candidate targets for drug repurposing in Alzheimer’s disease. Alzheimer’s research & therapy, 14(1), 7. 10.1186/s13195-021-00951-z

Kaul, G., Akhir, A., Shukla, M., Shafi, H., Akunuri, R., Pawar, G., Ghouse, M., Srinivas, N., & Chopra, S. (2023). Oxiconazole Potentiates Gentamicin against Gentamicin-Resistant Staphylococcus aureus *In Vitro* and *In Vivo*. Microbiology spectrum, 11(4), e0503122. 10.1128/spectrum.05031-22

Kaul, G., Akhir, A., Shukla, M., Rawat, K. S., Sharma, C. P., Sangu, K. G., Rode, H. B., Goel, A., & Chopra, S. (2022). Nitazoxanide potentiates linezolid against linezolid-resistant Staphylococcus aureus in vitro and in vivo. The Journal of antimicrobial chemotherapy, 77(9), 2456–2460. 10.1093/jac/dkac201

Knox, C., Wilson, M., Klinger, C. M., Franklin, M., Oler, E., Wilson, A., Pon, A., Cox, J., Chin, N. E. L., Strawbridge, S. A., Garcia-Patino, M., Kruger, R., Sivakumaran, A., Sanford, S., Doshi, R., Khetarpal, N., Fatokun, O., Doucet, D., Zubkowski, A., Rayat, D. Y., … Wishart, D. S. (2024). DrugBank 6.0: the DrugBank Knowledgebase for 2024. Nucleic acids research, 52(D1), D1265–D1275. 10.1093/nar/gkad976

Lee, A. S., de Lencastre, H., Garau, J., Kluytmans, J., Malhotra-Kumar, S., Peschel, A., & Harbarth, S. (2018). Methicillin-resistant Staphylococcus aureus. Nature Reviews Disease Primers, 4(1), 18033. 10.1038/nrdp.2018.33

Li, P., Wang, J., Qiao, Y., Chen, H., Yu, Y., Yao, X., Gao, P., Xie, G., & Song, S. (2021). An effective self-supervised framework for learning expressive molecular global representations to drug discovery. Briefings in Bioinformatics, 22(6). 10.1093/bib/bbab109

London A. J. (2019). Artificial Intelligence and Black-Box Medical Decisions: Accuracy versus Explainability. The Hastings Center report, 49(1), 15–21. 10.1002/hast.973

Muegge, I., & Mukherjee, P. (2015). An overview of molecular fingerprint similarity search in virtual screening. Expert Opinion on Drug Discovery, 11(2), 137–148. 10.1517/17460441.2016.1117070

Pedregosa, F., Varoquaux, G., Gramfort, A., Michel, V., Thirion, B., Grisel, O., Blondel, M., Prettenhofer, P., Weiss, R., Dubourg, V., Vanderplas, J., Passos, A., Cournapeau, D., Brucher, M., Perrot, M., & Duchesnay, E. (2011). Scikit-learn: Machine Learning in Python. Journal of Machine Learning Research, 12, 2825–2830.

Qiu, Y., & Cheng, F. (2024). Artificial intelligence for drug discovery and development in Alzheimer’s disease. Current opinion in structural biology, 85, 102776. 10.1016/j.sbi.2024.102776 RDKit: Open-source cheminformatics. https://www.rdkit.org

Selvaraju, R. R., Cogswell, M., Das, A., Vedantam, R., Parikh, D., & Batra, D. (2019). Grad-CAM: Visual Explanations from Deep Networks via Gradient-Based Localization. International Journal of Computer Vision, 128. 10.1007/s11263-019-01228-7

Stokes, J. M., Yang, K., Swanson, K., Jin, W., Cubillos-Ruiz, A., Donghia, N. M., MacNair, C. R., French, S., Carfrae, L. A., Bloom-Ackermann, Z., Tran, V. M., Chiappino-Pepe, A., Badran, A. H., Andrews, I. W., Chory, E. J., Church, G. M., Brown, E. D., Jaakkola, T. S., Barzilay, R., & Collins, J. J. (2020). A Deep Learning Approach to Antibiotic Discovery. Cell, 180(4), 688–702.e613. 10.1016/j.cell.2020.01.021

Wohlleben, W., Mast, Y., Stegmann, E., & Ziemert, N. (2016). Antibiotic drug discovery. Microbial biotechnology, 9(5), 541–548. 10.1111/1751-7915.12388

Wong, F., Zheng, E. J., Valeri, J. A., Donghia, N. M., Anahtar, M. N., Omori, S., Li, A., Cubillos-Ruiz, A., Krishnan, A., Jin, W., Manson, A. L., Friedrichs, J., Helbig, R., Hajian, B., Fiejtek, D. K., Wagner, F. F., Soutter, H. H., Earl, A. M., Stokes, J. M., . . . Collins, J. J. (2023). Discovery of a structural class of antibiotics with explainable deep learning. Nature, 626(7997), 177–185. 10.1038/s41586-023-06887-8

Zeng, X., Wang, F., Luo, Y., Kang, S. G., Tang, J., Lightstone, F. C., Fang, E. F., Cornell, W., Nussinov, R., & Cheng, F. (2022a). Deep generative molecular design reshapes drug discovery. *Cell reports*. Medicine, 3(12), 100794. 10.1016/j.xcrm.2022.100794

Zeng, X., Xiang, H., Yu, L., Wang, J., Li, K., Nussinov, R., & Cheng, F. (2022b). Accurate prediction of molecular properties and drug targets using a self-supervised image representation learning framework. Nature Machine Intelligence, 4(11), 1004–1016. 10.1038/s42256-022-00557-6

Zhou, Y., Hou, Y., Shen, J., Huang, Y., Martin, W., & Cheng, F. (2020a). Network-based drug repurposing for novel coronavirus 2019-nCoV/SARS-CoV-2. Cell Discovery, 6(1), 14. 10.1038/s41421-020-0153-3

Zhou, Y., Wang, F., Tang, J., Nussinov, R., & Cheng, F. (2020b). Artificial intelligence in COVID-19 drug repurposing. The Lancet Digital Health, 2(12). 10.1016/S2589-7500(20)30192-8

Zore, M., Gilbert-Girard, S., San-Martin-Galindo, P., Reigada, I., Hanski, L., Savijoki, K., Fallarero, A., Yli-Kauhaluoma, J., & Patel, J. Z. (2022). Repurposing the Sphingosine-1-Phosphate Receptor Modulator Etrasimod as an Antibacterial Agent Against Gram-Positive Bacteria. Frontiers in microbiology, 13, 926170. 10.3389/fmicb.2022.926170

